# Biological aging of two innate behaviors of *Drosophila melanogaster*: escape climbing versus courtship learning and memory

**DOI:** 10.1101/2023.10.10.561752

**Authors:** Jessica Thiem, Maria Viskadourou, Alexandros Gaitanidis, Dimitrios J Stravopodis, Roland Strauß, Carsten Duch, Christos Consoulas

## Abstract

Motor and cognitive aging can severely affect life quality of elderly people and burden health care systems. In search for diagnostic behavioral biomarkers, it has been suggested that walking speed can predict forms of cognitive decline, but in humans, it remains challenging to separate the effects of biological aging and lifestyle. We examined a possible association of motor and cognitive decline in Drosophila, a genetic model organism of healthy aging. Long term courtship memory is present in young flies but absent already during mid life (4-8 weeks). By contrast, courtship learning index and short term memory (STM) are surprisingly robust and remain stable through mid (4-8 weeks) and healthy late life (>8 weeks), until courtship performance collapses suddenly at ∼4.5 days prior to death. By contrast, climbing speed declines gradually during late life (>8 weeks). The collapse of courtship performance and short term memory close to the end of life are not related to the gradual late life decline in climbing speed. Thus, during healthy aging in Drosophila, climbing and courtship motor behaviors decline differentially, unlikely share a common cause, and motor and cognitive performance decline are not closely associated to each other.

## INTRODUCTION

In the face of increasing life expectancy in modern societies, aging-related decline in cognitive and motor performance becomes an increasing burden for health and social care systems (1,2). A fundamental prerequisite toward healthy aging is a better mechanistic understanding how the process of aging manifests in malfunction of the nervous system, but aging is a multi-causal, highly variable and individual process. In all animals, from worms (3) to flies (4) and humans (5,6), some individuals reach high ages without major impairments and are commonly referred to as wellderlies, whereas others suffer from prolonged periods of cognitive and/or motor decline. On the molecular level the nine main causes of aging are genomic instability, telomere attrition, epigenetic alterations, loss of proteostasis, deregulated nutrient sensing, mitochondrial dysfunction, cellular senescence, stem cell exhaustion, and altered intercellular communication (7). However, the resulting impairments on the nervous system and behavioral levels are diverse, and it is often difficult to separate the functional consequences of biological aging, diseases, and life-style. Some authors have proposed that one common cause underlies diverse aging phenotypes in the human nervous system, and this has been referred to as a unifying cause of aging (8). In support of this, positive correlations between specific aspects of motor function and cognitive decline have been provided (9,10) during human aging. In particular, in older adults, walking speed decrease is a good predictor of cognitive decline, but the reverse does not seem to be the case (11,12). On the other hand, ample evidence across species shows that different types of neurons and neural circuits (13,14), as well as different brain parts are differentially vulnerable to age-related and neurodegenerative decline (15). Within neurons, the synaptic compartment seems to be the most vulnerable one (15). Consequently, different cellular and circuit malfunctions have been attributed to underlie different age-related behavioral impairments. For example, in non-primate monkeys working memory decline has been related to high vulnerability of thin spines in prefrontal cortex (16), whereas loss of navigational ability has been attributed to reduced plasticity in place and grid cell circuitry (17). By contrast, the cellular causes for aging-related reduced locomotion speed that is apparent across species (9,18–20) remain largely unclear. Is there a unifying cause for multiple age-related impairments, and can some predict the occurrence of others? To start addressing this question, longitudinal studies of multiple aging-related nervous system malfunctions are needed (9).

As entry point toward addressing this question in the invertebrate genetic model system, *Drosophila melanogaster*, we investigate age-related decline of two well-studied innate behaviors in parallel, namely frustration learning during courtship (21) and escape climbing behavior (22). Drosophila has the advantage of a relatively short life-span of 60 to 90 days under laboratory conditions (4). First, this allows longitudinal studies within a few months. Second, neurodegenerative diseases are modeled in Drosophila upon introduction of human disease factors, but do not natively exist, so that longitudinal assessment can be conducted with healthy flies. This circumvents the problem that age-related changes in cognitive and physical functioning often result from interactions between the normal aging process and diseases with unknown onset, which makes their effects often indistinguishable (23). In addition, the genetic power of the Drosophila model system has advantaged to address the molecular mechanisms of aging-related cognitive and motor decline, and the relative simplicity of the nervous systems (∼100.000 neurons) may help identifying common or diverse cellular causes for cognitive and motor decline.

We have previously described the patterns of normal Drosophila motor aging and found that impairment onset, duration, and severity are highly variable and unpredictable (4), just as is the case in mammals. Moreover, flies can remain healthy until a few hours prior to death (wellderlies), or suffer from multiple days of combinations of different motor impairments (illderlies), again phenotyically similar to what has been reported for mammals (24,25), including humans (26). On the other hand, many studies have used Drosophila to investigate memory decline in old flies (27–31). However, knowledge on the relationship between the time course of motor and cognitive decline is sparse. We unravel the patterns of age-related decline in courtship learning and memory in parallel with that of locomotion decline to test whether one can predict the other, whether both occur in synchrony, or whether both manifest independently from each other.

## METHODS

### Animals, rearing conditions

Oregon-R (strain: # 5, Bloomington fly stock center) wild type flies were reared at 25°C and 70% relative humidity under a 12-h/12-h Light/Dark cycle. (rearing conditions:24±1 C°, 70% humidity; conditions during experimental trials:21±2 C° 60% humidity). Diet was based on corn flour-yeast agar medium containing 0.75% (w/v) agar, 4.5% (w/v) dry yeast, 3.5% (w/v) corn meal, 5.5% (w/v) Sucrose, 0.4% (v/v) Propionic acid, 2.5% (v/v) nipagen diluted in 10% absolute ethanol.

### Physical functioning and reactivity assay applied daily from 60 day of age until death

The Zeitgeber was a 12h light/dark cycle with light-on at 8am and light-off at 8pm with testing once/day at 10pm (light) in all males from the age of 60 days until death. Escape performance was tested in individual flies in their food vials by gently banging the vial on the counter. We applied a personalized assay described in (4). In brief in this assay a fly is aroused by gentle but abrupt tapings of the vial, that releases walking or/and climbing behaviors and thus allowing to assess the physical status and diagnose walking impairments. The resulting physiological status criteria are described in Figure 3 and in (4).

### Startle escape assay

Oregon-R males were singly isolated in falcon vials and left undisturbed for 30 minutes, enough time for acclimation in the new environment. Then each fly received six light stimuli (displacement of the fly at the bottom of the vial by tapping or flipping the vial) followed by six moderate stimuli (1 s vortexing) in two rounds. Finally, each fly received a combination of ∼20 mixed stimuli (flipping, banking and vortexing of the vial) in a fast pace. This fast delivery of different stimuli caused, at least one fierce escape response during which the fly performs close to its ability. We measured the time-to-climb a 6 cm vertical distance on the wall of the vial within 10 seconds. Failure was considered the unsuccessful effort of completing the task within 10 s.

### Courtship and courtship conditioning assay

Courtship assays were performed in mating chambers (10 mm diameter, 5 mm depth) and recorded with digital camcorders (Sony HDR-XR260). Flies were collected on the day of eclosion and kept in a 25°C incubator with a 12/12 hour L/D cycle. Naive males were kept in individual test tubes and target females and males for mating were store in small groups. The total amount of time a male was engaged in courtship activity with an unanesthetized target female (trainer or tester) during a test period of 10 min or until successful copulation occurred was scored. Overall, we followed the experimental procedures described in (32). For courtship condition assays, a single test male was placed in the chamber with a mated female (trainer) for one hour. The first and last 10 minutes were recorded and analyzed. After a 5 min isolation period, the trained male faced a tester (unanesthetized virgin female) and the 10 first minutes were recorded. A sham-trained male was kept alone in the courtship chamber for one hour and paired with a tester female for 10 min. Thus, for each 10-min recording, we calculated the courtship index (CI), CI initial, CI final, CI test, and CI sham. For long-term memory we followed the courtship conditioning assay protocol as described in (33). The learning index is the ratio of the courtship level in the final 10 min of the training (CI final) to that of the initial 10 min (CI initial). The memory index is the ratio CI test/the mean of CI sham. A memory index close to 1 indicates that there is no memory because the courtship level of the trained males is similar to that of the sham-trained males.

### Statistical analysis

All measurements were tested for Gaussian distribution by D’Agostino & Pearson omnibus normality test. When the data was following Gaussian distribution, the unpaired t-test was used. For comparison of two samples with data not following Gaussian distribution the two-tailed Mann–Whitney-U tests (95% confidence level, statistical significance P<0.05) was performed. For comparison of more than two groups, Kruskal–Wallis ANOVA and Dunn’s post-hoc test for multiple comparisons (statistical significance P<0.05) was used. To display the measurements, box-and-whisker plots were chosen, and medians were used as central values. Boxes included the medial 5–95%, and the whiskers included min to max of the data values. Outliers were not shown. Significant differences were accepted at p< 0.05. All statistical analyses in this study were performed using GraphPad Prism version 6.00 for Windows, (GraphPad Software, La Jolla, CA, USA).

## RESULTS

We used two well studied innate behaviors (escape climbing and courtship) to uncover the age-dependent changes in physical and cognitive functioning of male flies during mid life (4-8 weeks) and late life (>10 weeks). First, we examined the physical status of mid life and late life males in the escape climbing assay. Flies housed in cylindrical vials can be startled by mechanical stimuli, such as banging of the vial onto the top of the counter. In response to this disturbance the animal immediately escapes by climbing up the wall of the vial, obeying its innate negative geotaxis tendency. Less common is escape by jumping, or flight. Previous studies have demonstrated that climbing performance gradually decays with age (22), and that climbing ability is a good proxy for physiological age, whereas climbing impairment is a predictor of death (4).

In preliminary studies we noticed that escape climbing is scalable with the arousal level of the individual and the intensity of stimulation. Arousal level was controlled for as good as possible by treating all individuals identically for 30 minutes before testing (see methods). To account for stimulus intensity, we used three different stimulation regimes (see methods) for measuring escape performance (time to complete climbing a 6 cm vertical distance) in individual male flies of four different age groups (4, 6, 8, 10 weeks). We found that the strongest stimulation induced the fastest climbing responses across all ages tested (Fig. 1A). Independently of the type and intensity of the stimulation regime, climbing performance remains constant during mid life (4-8 weeks of age), but decreases significantly between the 8^th^ and 10^th^ week (Fig. 1A). Similarly, the percentage of successful responses (completion of the task within 10 s) drops from ∼80% in 8 weeks old flies to 20% −40% (depending on stimulus strength) in 10 weeks old flies (Fig. 1B). Thus, at a critical time after the eighth week of age, males become slower and fail more often to complete the task, but these flies are still capable of performing escape climbing.

**Figure 1.**
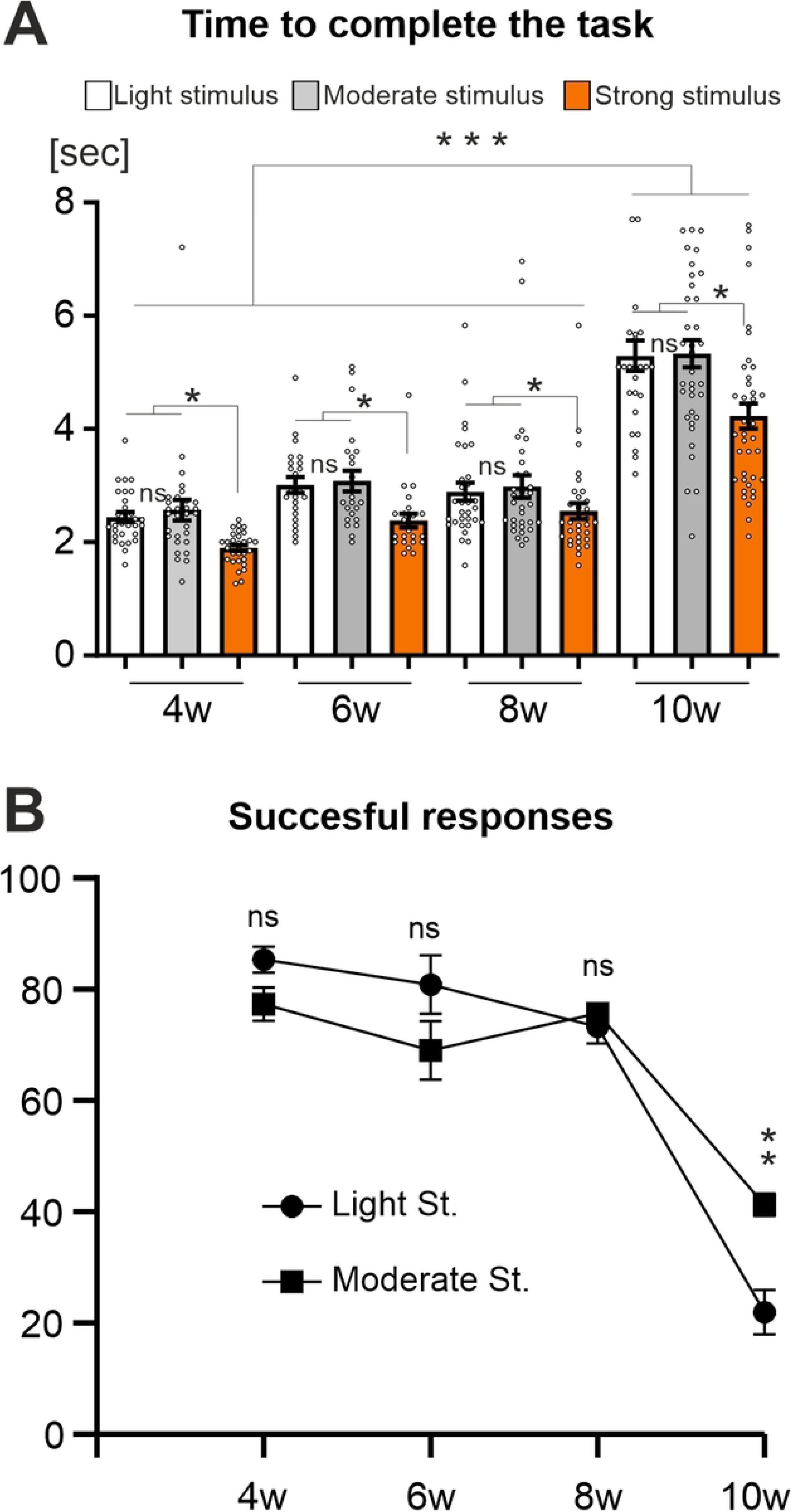
Climbing performance remains stable in mid life but decreases during late life (A) Male flies of different ages were individually tested for their performance in climbing a 6 cm vertical distance after having been startled with different stimulation regimes. The strongest stimuli caused the fastest responses at all ages. **(B)** Successful responses (completion of the 6 cm climbing task within 10 s) remain unaltered between 4 and 8 weeks of age (mid life) but decline after the 8^th^ week (late life). (One-Way ANOVA, Kruskal-Wallis multiple comparison test, * p<0.05, ** p<0.01, *** p<0.001).

Since we aim to unravel the patterns of age-dependent escape climbing decay and relate this to courtship learning and memory tasks, longitudinal testing of individual flies is necessary. Flies develop individual locomotor and behavioral disabilities prior to their natural death independently of the age they die (4). This pre-death morbidity period can vary from few hours to several days, has an unpredictable onset, and requires daily testing to be traced. Moreover, for measuring courtship learning and memory indexes daily, it is imperative to first investigate to what extent 24 hours long term memory (LTM) traces from the previous day affect the outcome of the next day’s test. We therefore estimated the 24 hours courtship LTM in males of 4, 8, and 10 weeks of age (Figure 2). All males were fit individuals capable of performing escape climbing.

**Figure 2.**
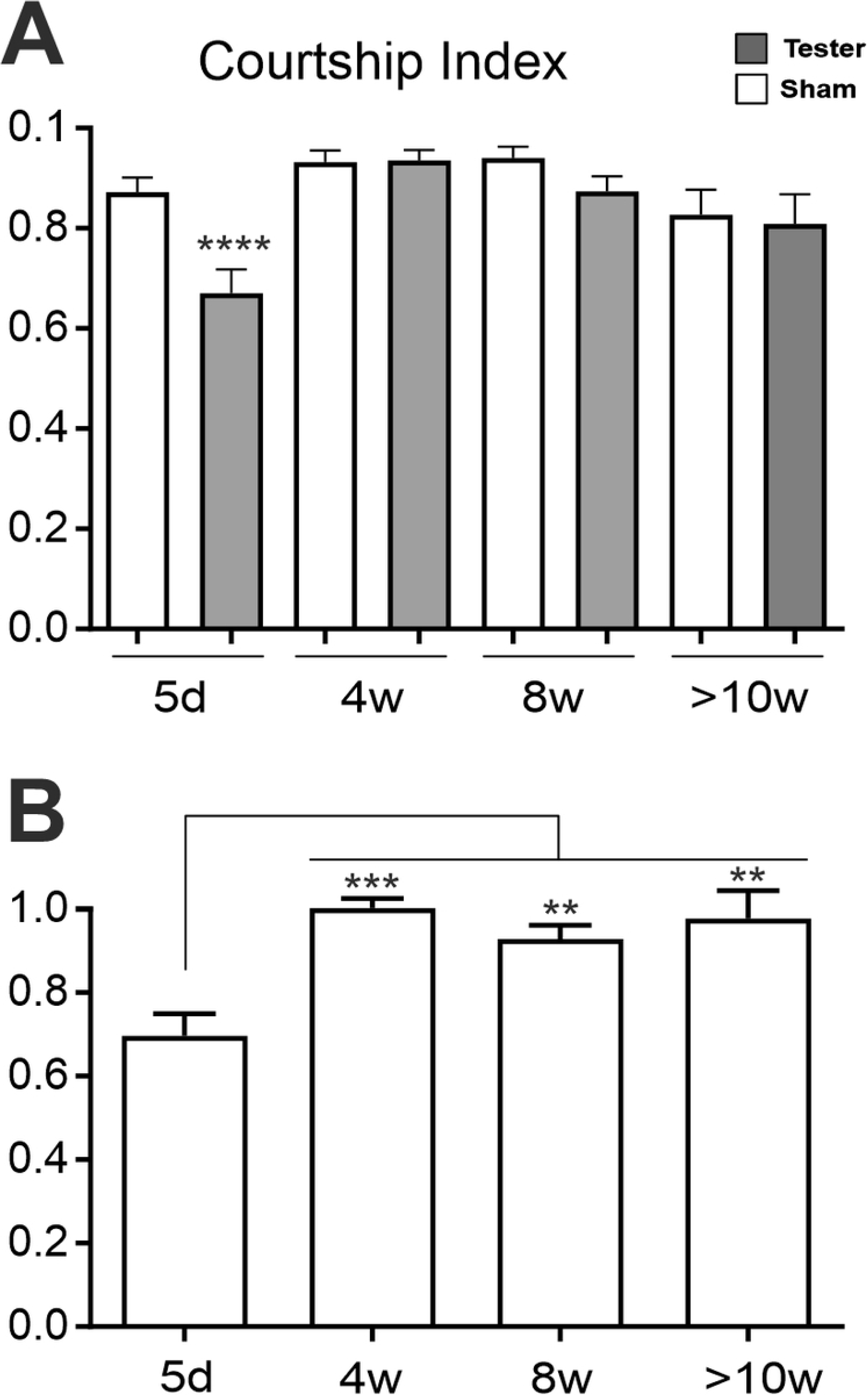
courtship LTM is absent during mid and late life (A) When presented with a virgin female, courtship indices of the Tester (males presented with a mated female on the previous day) and Sham males (naïve males) differ in young flies (5d), thus indicating courtship LTM. However, Tester and Sham courtship indices are not different during mid and late life. Courtship index was estimated as the percentage of male courtship within a 10 minutes time window. Tested males first courted towards a mated female for 8 hours, were then isolated for 24 hours, and finally courted against a virgin tester female. **(B)** LTM (24 hours) courtship memory index in mid and late life. Note that the LTM index approaches 1 at 4 weeks and all older ages tested, indicating absence of LTM from 4 weeks on. (T-test for A, One-Way Anova, Kruskal-Wallis multiple comparison test for B. ** p<0.01, *** p<0.001, **** p<0.0001).

Courtship index was similarly high in both, the tested males (confronted with a virgin female, but subjected to rejection by a mated female on the previous day) and the sham males (not previously frustrated, Fig. 2Α), and thus the long-term memory (LTM) index approaches 1 at all ages of 4 weeks and older (Fig. 2Β). This shows that no courtship memory is produced by this conditioning with a previously mated female target at any of the mid to advanced ages tested. However, courtship LTM is present in young flies (Fig. 2B), as previously shown by others (34). Ensuring that males do not remember their frustration with the mated female next day, the precondition for daily testing of courtship conditioning and escape climbing in longitudinal studies is fulfilled.

We next performed a longitudinal study with 53 males to study motor and cognitive performance, measured in climbing and courtship conditioning assays (Fig. 3). All animals for which daily fitness testing detected pre-death morbidity, such as an inability to climb, were not tested on subsequent days. Therefore, all animals were tested until they reached the pre-death impairment period (Figs. 3A, B). Twenty-five of these flies reached ages >10 weeks without impairment and were tested daily until pre-death morbidity. In the same experimental run, another population of single males (N=42) was not tested but scored for survival (control). Experimental testing did not reduce lifespan (Fig. 3C). An event history chart summarizes health-span (Fig. 3A, gray bars), impairment-span (Fig. 3A, B, colored bars), and lifespan for all 53 flies (Fig. 3A). Startle response impairments (color code for impairment type, Fig. 3D) were detected in 60.4% of all individuals on average 1.5 days prior to death (Fig. 3F). The remaining 39.6% were devoid of pathological phenotypes (Fig. 3A) and responded normally to the stimulation by climbing, jumping, or flying, until 1 day prior to death (Fig. 3A, B). As previously shown (4), even these “wellderly” flies eventually develop impairments for some hours prior to death. We measured the climbing speed of all fit males up to 11 weeks of age (Fig. 3E). Maximum and average climbing speed are lower in mid aged flies than reported for young flies (19), remain unaltered throughout mid life (4-8 weeks), but decline significantly between the 8th and 9th week (Fig. 3E). This pattern of decay confirmed our finding from the cross-sectional study (Fig. 1).

To relate escape climbing performance and courtship (performance, learning, and short-term memory) to each other, the same flies were also tested in the courtship conditioning assay (Fig. 4A). Males were presented for 1h to mated females (trainer female) and courtship index was measured during the first 10 minutes (CI initial, Fig. 4A, black bars) and the last 10 minutes (CI final, Fig. 4A, gray bars). The ratio of CI final/CI initial is the learning index during training, which shows no statistical differences between 4 and 10 weeks of age (Fig. 4B). Next, 5 minutes after the 1h training, the trained male was presented to a young virgin female (CI test, Fig. 4A, dark gray bars). In parallel, same aged, non-trained males were presented to a young virgin female (CI Sham, Fig.4A, white bars). The ratio of CI test divided by CI Sham yields the 5 minutes short term memory (STM) index, which also shows no statistically significant differences between 4 and 10 weeks of age (Fig. 4C).

**Figure 3.**
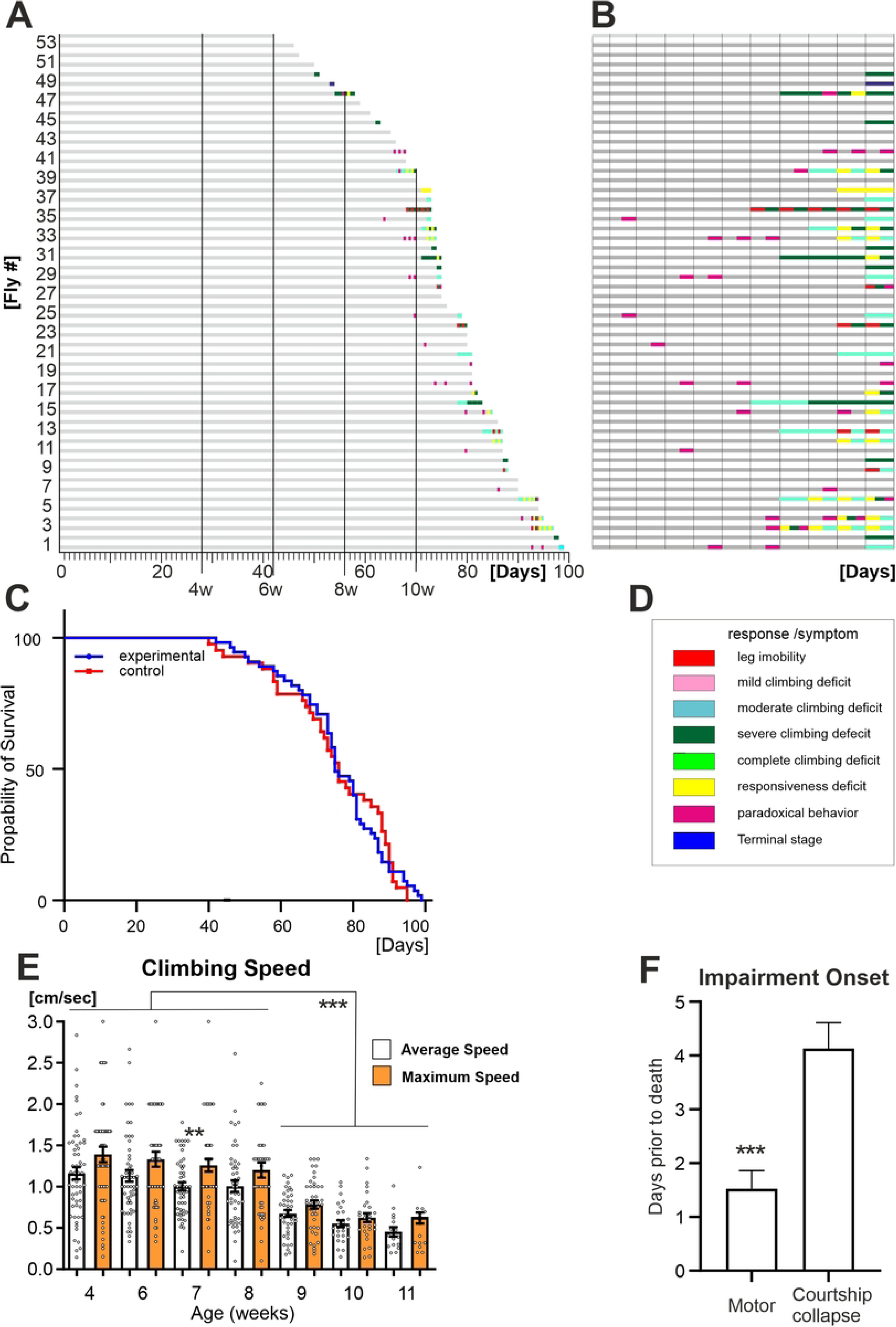
Late life pathophysiology of locomotor behavior (A) Life history chart for 53 male Oregon-R flies tested daily and individually in the startle assay from the age of 60 days until death. Gray bars indicate health-span and colored bars disabilities of different categories. 39.6% (21 out of 53) of the flies show no sign of impairment until the last day of life. **(B)** Time enlargement of the last 10 days for all flies. **(C)** Survival curves for the tested population in **A** and non-tested control flies are not significantly different [Log-rank (Mantel-Cox) test; X^2^= 0.001431, p= 0.9698]. (**D**) Color-code for behavioral impairments as defined in Gaitanidis et al. (2019). (**E**) Maximum climbing speed drops significantly between the 8^th^ and 9^th^ week. (**F**) Severe reduction (>90%) in male courtship occurs on average 4 days prior to death, 2.5 days earlier than the onset of motor impairments. (One-Way ANOVA, Kruskal-Wallis multiple comparison test for E; T-test for F. ** p<0.01, *** p<0.001).

**Figure 4.**
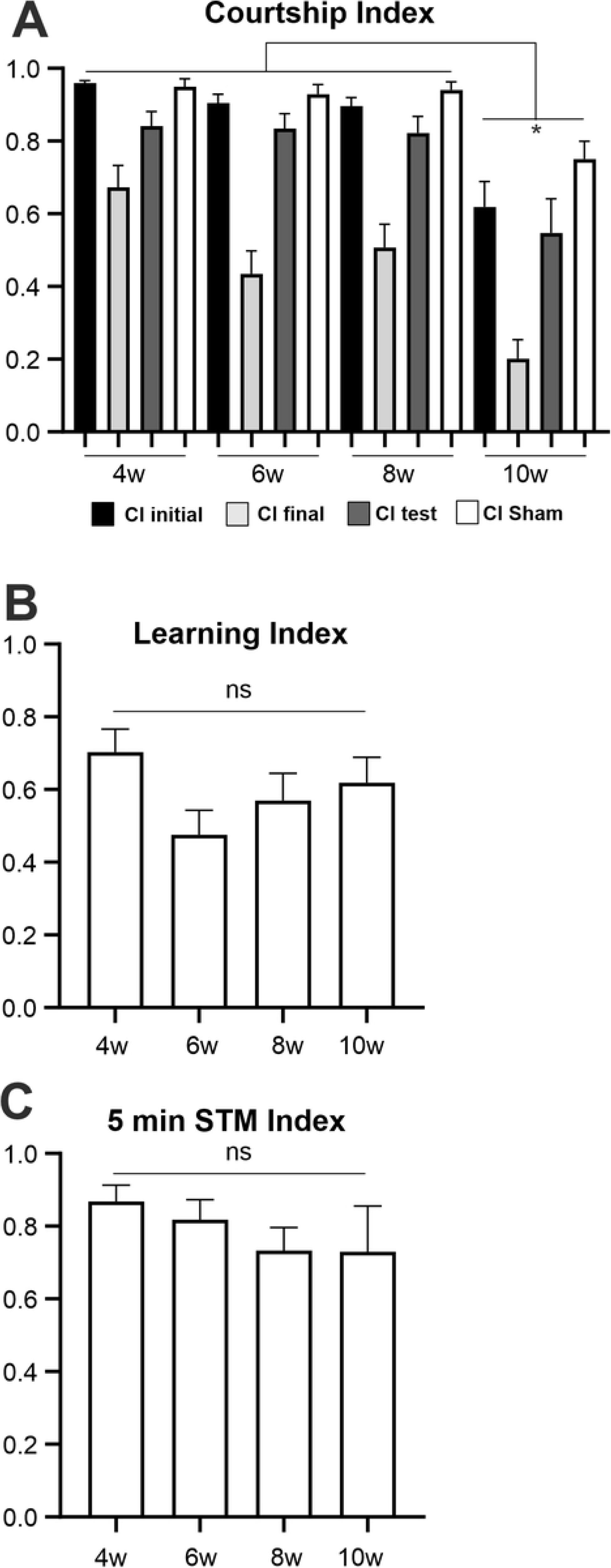
Male courtship index decays in old flies, but courtship learning and (STM) remain stable. (**A**). Courtship Index is estimated as the percentage of male courtship within a 10 minute time window. Courtship conditioning assays were performed with the males of different ages (4, 6, 8, and 10 weeks of age). First the trained male courts against a mated female trainer for one hour [recorded:10 first min with trainer (CI initial), 10 last mins with trainer (CI final)]. Next, after 5 min isolation the trained male is presented to a young virgin female tester [recorded:10 first mins with the virgin tester (CI test)]. As a control, age-matched, non-trained males (Sham) are presented with a virgin female. (**B**) The learning index is the ratio of the courtship indices during the final 10 min of the training (CI final) and that of the initial 10 min (CI initial). (**C**) The memory index is calculated by dividing CI test by the mean of the sham control courtship levels (CI sham).

Courtship motor performance is robust during all ages, CI initial and CI Sham are constantly high (>0.9) between 4 and 8 weeks, but decay after the 8th week of age (Fig. 4A). Further analysis of courtship (orientation), singing, licking and attempting copulation latencies (the time between the introduction of flies in the courtship chamber to the time of the first exhibition of any of the particular behaviors) reveal a gradual age-dependent increase in the latency of male engagement towards all three female partners (S1 Fig, Table 1). However, at any given age tested no differences are found in the latencies of engagement with the mated trained, the virgin tester, or the virgin control females (S1 Fig, and see below). The age-related latency increase accumulates to a decrease of CI in 10 weeks old flies (Fig. 4A). Learning index during training varies between 0.45 to 0.70 with no significant differences between age groups and no clear trend over time (Fig. 4B), indicating persistence of learning ability throughout ages. The STM memory index varies between 0.7 to 0.85 (Fig. 4C) without statistically significant differences between ages. Therefore, in animals without major physical impairments both courtship learning and STM are stable throughout life. In contrast, climbing and courtship motor performance is robust in mid life up to the age of 8 weeks but declines at old ages (8-10 weeks), although 10 weeks old, wellderly flies can still perform.

**Table 1.** Comparison of latencies for male courtship parameters depicted in Figure 4B. p -values from One-way ANOVA, Kruskal-Wallis test, Tukey’s multiple comparisons test. Values in bold indicate significance.

|  | Courtship<br>Latency | Singing<br>Latency | Licking<br>Latency | Att.<br>Cop.<br>Latency |
| --- | --- | --- | --- | --- |
| 4W Tester vs. 4W<br>Trainer | >0.9999 | >0.9999 | >0.9999 | 0.956 |
| 4W Tester vs. 4W<br>Sham | >0.9999 | >0.9999 | >0.9999 | >0.9999 |
| 4W Trainer vs. 4W<br>Sham | >0.9999 | >0.9999 | >0.9999 | >0.9999 |
| 6W Tester vs. 6W<br>Trainer | >0.9999 | >0.9999 | >0.9999 | 0.9934 |
| 6W Tester vs. 6W<br>Sham | >0.9999 | >0.9999 | >0.9999 | 0.0031 |
| 6W Trainer vs. 6W<br>Sham | >0.9999 | >0.9999 | >0.9999 | **<0.0001** |
| 8W Tester vs. 8W<br>Trainer | >0.9999 | >0.9999 | >0.9999 | 0.9999 |
| 8W Tester vs. 8W<br>Sham | 0.4614 | >0.9999 | >0.9999 | 0.2567 |
| 8W Trainer vs. 8W<br>Sham | 0.0039 | 0.0231 | <0.0001 | 0.5728 |
| 10W Tester vs. 10W<br>Trainer | >0.9999 | >0.9999 | >0.9999 | 0.9539 |
| 10W Tester vs. 10W<br>Sham | >0.9999 | >0.9999 | >0.9999 | 0.7747 |
| 10W Trainer vs. 10W<br>Sham | >0.9999 | >0.9999 | >0.9999 | >0.9999 |
| 4W Tester vs. 6W<br>Tester | 0.468 | 0.3145 | **0.0011** | **0.0026** |
| 4W Tester vs. 8W<br>Tester | 0.121 | **0.0433** | **<0.0001** | **0.0001** |
| 4W Tester vs. 10W<br>Tester | 0.0591 | 0.1366 | **<0.0001** | **0.0355** |
| 6W Tester vs. 8W<br>Tester | >0.9999 | >0.9999 | >0.9999 | >0.9999 |
| 6W Tester vs. 10W<br>Tester | >0.9999 | >0.9999 | 0.2871 | >0.9999 |
| 8W Tester vs. 10W<br>Tester | >0.9999 | >0.9999 | >0.9999 | >0.9999 |
| 4W Trainer vs. 6W<br>Trainer | 0.7131 | **<0.0001** | **0.0524** | **0.0013** |
| 4W Trainer vs. 8W<br>Trainer | **0.0015** | **0.001** | **<0.0001** | **<0.0001** |
| 4W Trainer vs. 10W<br>Trainer | **0.0134** | **0.0003** | **<0.0001** | **<0.0001** |
| 6W Trainer vs. 8W<br>Trainer | 0.2115 | **0.0518** | **0.0005** | **0.0098** |
| 6W Trainer vs. 10W<br>Trainer | 0.4979 | >0.9999 | **<0.0001** | **0.0846** |
| 8W Trainer vs. 10W<br>Trainer | >0.9999 | >0.9999 | **0.025** | >0.9999 |
| 4W Sham vs. 6W<br>Sham | 0.4722 | 0.2239 | **0.0155** | **<0.0001** |
| 4W Sham vs. 8W<br>Sham | >0.9999 | 0.0002 | **<0.0001** | **0.0004** |
| 4W Sham vs. 10W<br>Sham | 0.0131 | 0.0003 | **<0.0001** | **<0.0001** |
| 6W Sham vs. 8W<br>Sham | >0.9999 | 0.2272 | 0.1476 | >0.9999 |
| 6W Sham vs. 10W<br>Sham | 0.8516 | 0.0998 | 0.0739 | >0.9999 |
| 8W Sham vs. 10W<br>Sham | 0.2314 | >0.9999 | >0.9999 | >0.9999 |

To further define the patterns of decay in climbing speed and courtship performance in highly advanced ages, we conducted daily assays in twenty-five old (>10 weeks) wellderly males until they developed pre-death impairments or suddenly died (Fig. 3A, Fly # 1-25). The final decrease of climbing speed of elderly males occurs past the 75^th^ day of the cohort’s age (Fig. 5A) but CI and STM do not change during this time (Fig. 5B). Latencies in the initiation of courtship rituals remain stable throughout this period as well, with shorter latencies in courtship engagements towards the tester (virgin female; S2 Fig). Independent of the age at death, high levels of courtship performance against virgin and mated female partners drop suddenly, within one day (Fig. 5C) to negligible levels (CI<0.1). Therefore, on average courtship collapses occurs2.5 days prior to impairment onset and ∼4 days prior to death (Fig. 3F). By contrast, climbing speed declines gradually during the time when courtship performance collapses in the same individuals (Fig. 5C). One possibility to explain how gradual decreases in climbing speed through late age could cause a sudden collapse of courtship within one day would be a threshold for climbing/walking speed below which animals become incapable to court. However, this is not the case. First, plotting climbing speed versus CI, STM and the latency in the initiation of each courtship sub-behavior against the trainer (mated female) and tester (virgin female) for each individual male reveals either no correlation (ages: 4 weeks-10; Table 2) or week correlation (age: >10 weeks; Table 3). Second, plotting speed versus CI for old flies shows that some of the slowest males acquired high scores in in courtship performance and *vice versa*, other flies showed low CI but still had medium walking/climbing speed (Fig. 5D) These data indicate that the sudden drop in courtship performance at late ages is mechanistically independent of the gradual agerelated decline in locomotion speed.

**Figure 5.**
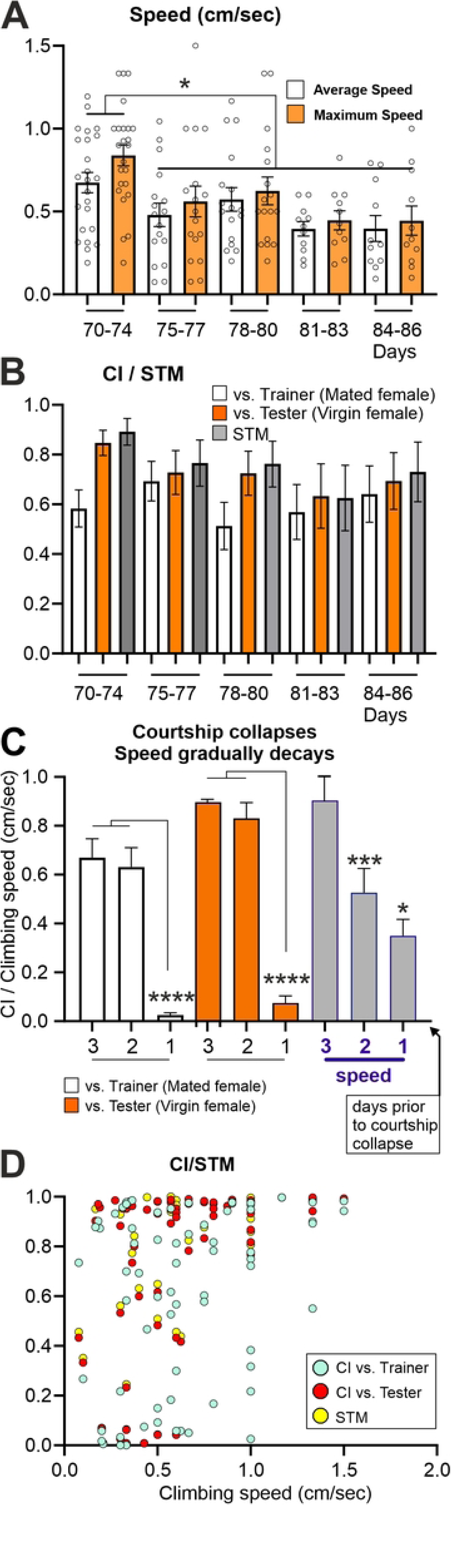
In old flies, speed decays gradually whereas courtship performances collapses, but a weak-to-moderate correlation is found between speed and STM. (A) Climbing speed measurements during the last days of the cohort (>10 weeks). Not the final drop of speed past the 75^th^ day. **(B)** Courtship and Short Term Memory (STM) Indexes remain stable during the last days. (**C**) Patterns of change in courtship and climbing speed during the last three days prior to courtship collapse. Note that courtship against the tester (CI test) and the trainer (CI initial) abruptly drop to very low levels whereas the climbing speed declines gradually. (One-Way ANOVA, Kruskal-Wallis multiple comparison test, * p<0.05, ** p<0.01, *** p<0.001, **** p<0.0001). **(D)** No correlation between speed and both courtship (CI trainer, CI tester) and STM indexes. Note that a slow individual can out-perform in courtship engagement and vice versa.

**Table 2.** 4-10 week old males. R and p values from Spearman rank correlation test. Values in bold indicate significance.

| Age | Speed vs. CI<br>initial | Speed vs. CI<br>test/STM | Speed vs.<br>Learning Index |
| --- | --- | --- | --- |
| 4 weeks | r=-0.09049<br>p= 0.5193 | r=-0.06935<br>p=0.6287 | r=-0.3567<br>p=0.0204 |
| 6 weeks | r=0.0266<br>p=0.869 | r=0.2611<br>p=0.070 | r=0.2333<br>p=0.147 |
| 8 weeks | r=0.1523<br>p=0.361 | r=0.1089<br>p=0.492 | r=-0.0126<br>p=0.942 |
| 10 weeks | **r=0.5042**<br>**p=0.019** | r=0.3515<br>p=0.129 | r=-0.01355<br>p=0.955 |

**Table 3.** >10 week old males. R and p values from Spearman rank correlation test. Values in bold indicate significance.

|  | **Towards trainer (Initial 10 mins)** |  |  |  |  |
| --- | --- | --- | --- | --- | --- |
| r, p /<br>Age:<br>70-85 d | Speed<br>vs.<br>Orientation<br>Latency | Speed<br>vs.<br>Singing<br>Latency | Speed<br>vs.<br>Licking<br>Latency | Speed<br>vs.<br>At. Copulation<br>Latency | Speed<br>vs.<br>CI Initial |
| **r** | -0,1059 | -0,2100 | -0,2318 | -0,1805 | 0,3059 |
| **P** | 0,3831 | 0,1238 | 0,1129 | 0,2353 | **0,0085** |
|  | **Towards tester (Initial 10 mins)** |  |  |  |  |
| r, p /<br>Age:<br>70-85 d | Speed<br>vs.<br>Orient.<br>Latency | Speed<br>vs.<br>Singing<br>Latency | Speed<br>vs.<br>Licking<br>Latency | Speed<br>vs.<br>At. Copulation<br>Latency | Speed<br>vs.<br>CI test./<br>STM |
| **r** | -0,4022 | -0,4171 | -0,3212 | -0,3322 | 0,5169 |
| **P** | **0,0004** | **0,0005** | **0,0202** | **0,0184** | **<0,0001** |

Importantly, the orientation of the male towards the female, which is the first step in the escalating sequence of courtship rituals (measured as courtship/orientation latency, S1 Fig, S2 Fig), is largely independent of climbing performance across ages, including very old wellderly flies (Tables 2, 3) and even impaired individuals (our observation).

This suggests that both courtship target identification by the male and the innate interest to court are abilities that persists until almost death. Overall, the data indicate that during biological aging of otherwise healthy flies, age-related cognitive and motor decline occur with different dynamics and largely independently from each other (see discussion).

## Discussion

We used two robust innate behaviors with high evolutionary relevance for crosssectional and longitudinal assessment of aging-related cognitive and motor decline. Adequate courtship behavior ensures reproductive success and escape climbing is important for survival. Below we will first discuss the impact of aging on escape climbing, second age-related changes in courtship performance as well as in courtship learning and memory, and third, the relationship between cognitive and motor decline.

### The patterns of age-related escape climbing speed decrease

Startle induced escape responses can be elicited until few hours prior to death, unless flies suffer from locomotion impairments that make the task impossible. Such severely impaired flies were excluded from the analysis because we aimed at identifying the patterns of healthy motor aging. It is known that locomotion speed declines with age across species, including humans (35, 36) and Drosophila (19). Thus, age-related reductions in locomotion speed are a conserved feature of aging from flies to humans. However, both our cross-sectional as well as our longitudinal analyses show that once flies reach mid age (4 weeks), locomotion speed remains stable until the age of 8 weeks, but thereafter declines gradually again until late life (8-11 weeks). The same is the case for the responsiveness to the startle stimulus. Therefore, mid life is characterized by a locomotion performance plateau. This mid age-locomotor performance plateau in Drosophila is not in agreement with a previous report (19), but there, the Oregon-R wildtype strain used in our analysis has been tested only until 6 weeks of age, which might have hidden the performance plateau from the analysis. Similar observations have been made in human subjects. Although human physical performance often declines during mid age, assessment of marathon and half-marathon runners aged 20 to 79 years revealed no physical performance decline until the age of 55 years in subjects with healthy life-style. Parallel assessment with life-style questionaries has yielded the conclusion that human mid age physical performance decline is mainly attributed to life-style but not to biological aging (37). In our analysis on Drosophila, lifestyle was not a factor, because all animals were raised under identical laboratory conditions. As mentioned above, during late life, climbing speed and thus locomotor performance declines gradually both in Drosophila (8-11 weeks/56-77 days, this study) as well as in humans (55 to 79 years; (37).

### The patterns of courtship motor performance aging

Courtship behavior comprises a complex sequence of different motor tasks (orientation towards the female, “love-song” production by wing movement, chasing, licking, attempting copulation) which are orchestrated by communication between female and male flies (38, 21). The courtship index (CI) resembles the percentage of time during which any of these motor tasks is executed and was assessed during 10 minutes time periods in this study. Therefore, in contrast to escape climbing, which is a single motor task that is completed within seconds, CI rather represents a measure for perseverance of a complex motor behavior. Our longitudinal analysis of CI, and thus courtship motor performance, revealed a similar pattern of age-related decline as escape climbing speed. CI remains stable during mid life (four to eight weeks of age) but then declines significantly in old flies (>10 weeks). A similar tendency is apparent in the cross-sectional analysis, but this is not statistically significant, underscoring the importance of longitudinal assessment of individuals. These data indicate that both locomotor speed and motor behavioral perseverance are stable during mid life but start declining at similar old ages. Interestingly, the latencies to the first execution of the different parts of the courtship behavioral sequence remain unaltered throughout mid and late life, indicating that the initial interest of male flies to court remains unaltered, even in the face of declining motor abilities. In fact, even physically impaired males that can barely walk, but not sing, and approach death within ∼2 days show immediate orientation toward a virgin female, underscoring the robustness of innate male reproductive interest and target identification against biological aging.

During late life climbing speed decreases gradually, whereas courtship behavior collapses within one day. An obvious interpretation is that the gradual locomotion speed decline in old flies causes the courtship sequence to be interrupted, because the male can simply not chase the young, virgin female anymore. However, although there is a low-to-moderate correlation between walking speed and CI in old flies (>10 weeks), some of the slowest flies still show high CIs, and some reasonably fast flies show CIs close to zero. Therefore, the sudden collapse of courtship performance in old flies is unlikely a consequence of slow walking speed, but the patterns of late life decline differ for locomotion speed and motor perseverance.

### The patterns of courtship learning and memory aging

Although courtship is an innate behavior, its execution can be modified by learning. If courting males are rejected by a mated female, they subsequently engage less into courtship behavior when presented with a virgin female (39). In young male flies this memory can be retrieved either as STM shortly after the training with a mated female (here after 5 minutes) or as LTM 24 h after training (40). Knowledge on age-related changes of courtship memory is sparse, but it has been shown that courtship learning and memory assessed at 1h after training exists in mid aged flies (45 day of age), though reduced as compared to young animals (5 days of age, 41Sean et al., 2010). We assessed courtship learning and memory in parallel with motor aging during mid and late life of Drosophila. First, LTM as assessed 24hrs after training is absent already in mid aged flies. Similarly, specific olfactory memories are significantly reduced or absent in mid aged flies. Aversive olfactory memory acquired in a single olfactory conditioning session comprises three phases: short-term memory (STM), midterm memory (MTM), and longer-lasting anesthesia-resistant memory (ARM). Aging-related olfactory memory loss has been attributed to the decline of the amnesiac-dependent MTM (42, 27, 28), but LTM is also impaired (30). We find an age correspondence between the loss of olfactory MTM/LTM and courtship LTM in mid aged flies. By contrast, other LTMs, such as a body size memory that is formed during a critical period in young adults remains stable through all ages tested (43). In sum, in Drosophila different types of LTM can show different patterns of age-related decline. The absence of courtship LTM already after four weeks of adult life enabled us to conduct longitudinal assessment of learning and STM in the courtship assay during mid and late life, because we could exclude the interference of LTM with the repeated measurements throughout life from the same individuals.

In contrast to courtship LTM, learning during courtship as well as STM remain unaffected in late life until courtship behavior collapses entirely. This seems biologically relevant, because not learning that an ongoing or just waved courtship attempt is or has been futile would cause useless investment into a goal that cannot be achieved. In fact, recent data on olfactory memory indicate that the formation or retention of appetitive memory with survival benefits is maintained in aged flies (28). These data indicate that mechanisms that protect against cognitive loss may evolve predominantly for memories that are biologically meaningful also at late ages.

### Motor and cognitive decline show only moderate correlations and only during late life

Based on reported positive correlations between specific aspects of mobility and cognitive decline in humans (9–12) during aging, we originally hypothesized that age-related decreases in climbing speed may predict decline of courtship learning and memory defects. Our data largely reject this hypothesis. First, although climbing speed is slower in mid aged flies than in young flies, this cannot predict cognitive decline. Second, during mid life (4 to 8 weeks of age) we found no period of time during which motor decline can predict any aspect of cognitive decline in courtship learning and memory, or *vice versa*. LTM is already gone and STM remains unaltered through mid life. Only during late life (flies > 10 weeks) there is a moderate correlation between climbing speed and STM, but not between climbing speed and learning. Therefore, a common underlying cause for the different features of age-related nervous system decline that were investigated in this study seems unlikely. By contrast, the data favor the idea that various aspects of motor and cognitive decline are affected differentially by the aging process. This is in line with differential vulnerabilities of different types of neurons and neural circuits (13, 14), as reported across species. In Drosophila, genes and neural circuits underling escape motor responses (44) and courtship (45) are well characterized. Combining the recent success in circuit mapping in the relatively simple Drosophila brain, at least when compared to humans, with the versatile tools for genetic manipulation with exquisite spatial and temporal resolution makes Drosophila a useful genetic model to probe the molecular and cellular mechanisms underlying differential decline of distinct aspects of motor and cognitive function during biological aging.

---

**S1 Fig.**
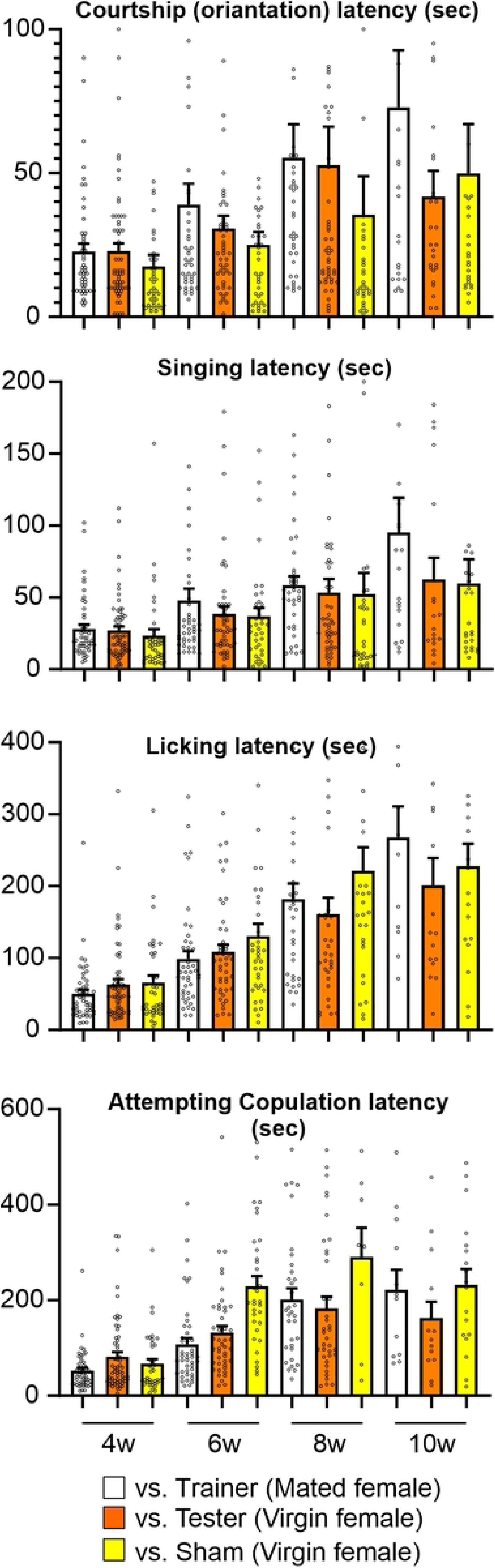
Courtship rituals latencies, that is the time it takes for an individual to initiate any of the behavior, increase in late life.

**S2 Fig.**
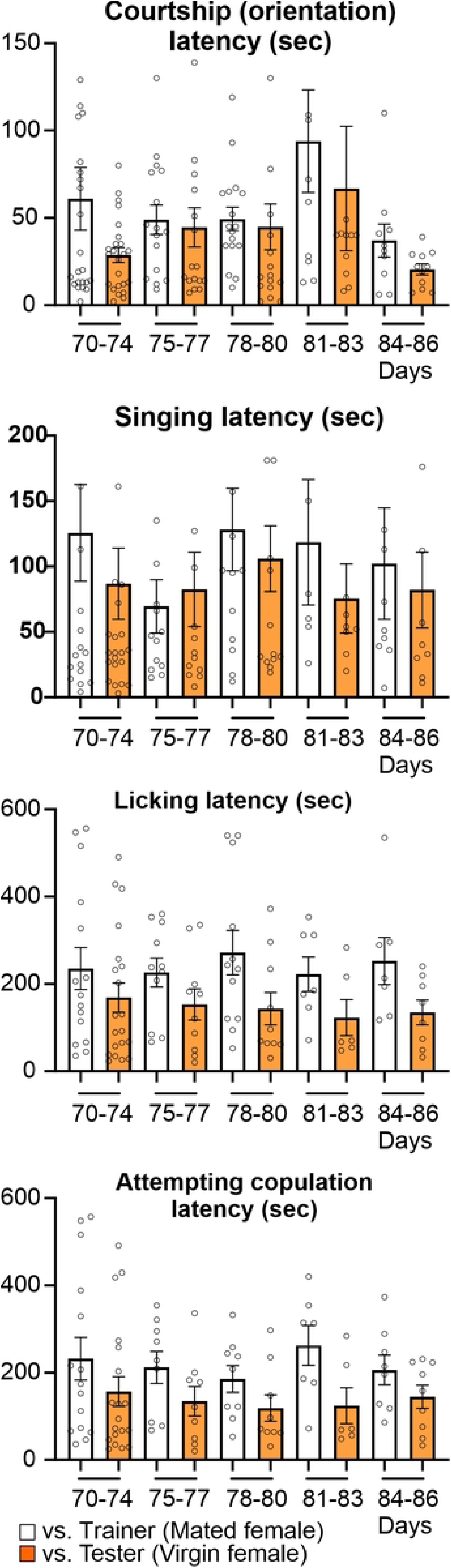
Latencies in the initiation of courtship rituals remain stable in old flies (>10 weeks).

## Notes

### Competing Interest Statement

The authors have declared no competing interest.

